# The initial gut microbiota and response to antibiotic perturbation influence *Clostridioides difficile* colonization in mice

**DOI:** 10.1101/2020.08.27.271304

**Authors:** Sarah Tomkovich, Joshua M.A. Stough, Lucas Bishop, Patrick D. Schloss

## Abstract

The gut microbiota has a key role in determining susceptibility to *Clostridioides difficile* infections (CDIs). However, much of the mechanistic work examining CDIs in mouse models use animals obtained from a single source. We treated mice from 6 sources (2 University of Michigan colonies and 4 commercial vendors) with clindamycin, followed by a *C. difficile* challenge and then measured *C. difficile* colonization levels throughout the infection. The microbiota were profiled via 16S rRNA gene sequencing to examine the variation across sources and alterations due to clindamycin treatment and *C. difficile* challenge. While all mice were colonized 1-day post-infection, variation emerged from days 3-7 post-infection with animals from some sources colonized with *C. difficile* for longer and at higher levels. We identified bacteria that varied in relative abundance across sources and throughout the experiment. Some bacteria were consistently impacted by clindamycin treatment in all sources of mice including *Lachnospiraceae, Ruminococcaceae*, and *Enterobacteriaceae*. To identify bacteria that were most important to colonization regardless of the source, we created logistic regression models that successfully classified mice based on whether they cleared *C. difficile* by 7 days post-infection using community composition data at baseline, post-clindamycin, and 1-day post-infection. With these models, we identified 4 bacteria that were predictive of whether *C. difficile* cleared. They varied across sources (*Bacteroides*), were altered by clindamycin (*Porphyromonadaceae*), or both (*Enterobacteriaceae* and *Enterococcus*). Allowing for microbiota variation across sources better emulates human inter-individual variation and can help identify bacterial drivers of phenotypic variation in the context of CDIs.

**Importance:** *Clostridioides difficile* is a leading nosocomial infection. Although perturbation to the gut microbiota is an established risk, there is variation in who becomes asymptomatically colonized, develops an infection, or has adverse infection outcomes. Mouse models of *C. difficile* infection (CDI) are widely used to answer a variety of *C. difficile* pathogenesis questions. However, the inter-individual variation between mice from the same breeding facility is less than what is observed in humans. Therefore, we challenged mice from 6 different breeding colonies with *C. difficile*. We found that the starting microbial community structures and *C. difficile* persistence varied by the source of mice. Interestingly, a subset of the bacteria that varied across sources were associated with how long *C. difficile* was able to colonize. By increasing the inter-individual diversity of the starting communities, we were able to better model human diversity. This provided a more nuanced perspective of *C. difficile* pathogenesis.

## Introduction

Antibiotics are a common risk factor for *Clostridioides difficile* infections (CDIs) due to their effect on the intestinal microbiota, but there is variation in who goes on to develop severe or recurrent CDIs after exposure (1, 2). Additionally, asymptomatic colonization, where *C. difficile* is detectable, but symptoms are absent, has been documented in infants and adults (3, 4). The intestinal microbiota has been implicated in asymptomatic colonization (5, 6), susceptibility to CDIs (7), and adverse CDI outcomes (9–12). However, it is not clear how much inter-individual microbiota variation contributes to the range of outcomes observed after *C. difficile* exposure relative to other risk factors.

Mouse models of CDIs have been a great tool for understanding *C. difficile* pathogenesis (13). The number of CDI mouse model studies has grown substantially since Chen et al. published their C57BL/6 model in 2008, which disrupted the gut microbiota with antibiotics to enable *C. difficile* colonization and symptoms such as diarrhea and weight loss (14). CDI mouse models have been used to examine translationally relevant questions regarding *C. difficile*, including the role of the microbiota and efficacy of potential therapeutics for treating CDIs (15). However, variation in the microbiota between mice from the same breeding colony is much less than the inter-individual variation observed between humans (16, 17). Studying CDIs in mice with a homogeneous microbiota is likely to overstate the importance of individual mechanisms. Using mice that have a more heterogeneous microbiota would allow researchers to identify and validate more generalizable mechanisms responsible for CDI.

In the past, our group has attempted to introduce more variation into the mouse microbiota by using a variety of antibiotic treatments (18–21). An alternative approach to maximize microbiota variation is to use mice from multiple sources (22, 23). The differences between the microbiota of mice from vendors have been well documented and shown to influence susceptibility to a variety of diseases (24, 25), including enteric infections (22, 23, 26–30). Different research groups have also observed different CDI outcomes despite using similar murine models (13, 18, 21, 31–33). Here we examined how variation in the baseline microbiota and responses to clindamycin treatment in C57BL/6 mice from six different sources influenced susceptibility to *C. difficile* colonization and the time needed to clear the infection.

## Results

### The variation in the microbiota is high between mice from different sources

We obtained C57BL/6 mice from 6 different sources: two colonies from the University of Michigan that were split from each other in 2010 (the Young and Schloss lab colonies) and four commercial vendors: the Jackson Laboratory, Charles River Laboratories, Taconic Biosciences, and Envigo (which was formerly Harlan). These 4 vendors were chosen because they are commonly used for murine CDI studies (26, 34–40). Two experiments were conducted, approximately 3 months apart.

We sequenced the 16S rRNA gene from fecal samples collected from these mice after they acclimated to the University of Michigan animal housing environment. We first examined the alpha diversity across the 6 sources of mice. There was a significant difference in the richness (i.e. number of observed operational taxonomic units (OTUs)), but not Shannon diversity index across the sources of mice (*P*_FDR_ = 0.03 and *P*_FDR_ = 0.052, respectively; Fig. 1A-B and Tables S1-2). Next, we compared the community structure of mice (Fig. 1C). The source of mice and the interactions between the source and cage effects explained most of the observed variation between fecal communities (PERMANOVA combined R^2^ = 0.90, *P* < 0.001; Fig. 1C and Table S3). Mice that are co-housed tend to have similar gut microbiotas due to coprophagy (41). Since mice within the same source were housed together, it was not surprising that the cage effect also contributed to the observed community variation. There were some differences between the 2 experiments we conducted, as the experiment and cage effects significantly explained the observed community variation for the Schloss and Young lab mouse colonies (Fig. S2A-B and Table S4). However, most of the vendors also clustered by experiment (Fig. S2C-D, F), suggesting there was some community variation between the 2 experiments within each source, particularly for Schloss, Young, and Envigo mice (Fig. S2G-H). After finding differences at the community level, we next identified the bacteria that varied between sources of mice. There were 268 OTUs with relative abundances that were significantly different between the sources (Fig. 1D and Table S5). Though we saw differences between experiments at the community level, there were no OTUs that were significantly different between experiments within Schloss, Young, and Envigo mice (all *P* > 0.05). By using mice from six sources we were able to increase the variation in the starting communities to evaluate in a clindamycin-based CDI model.

**Figure 1.**
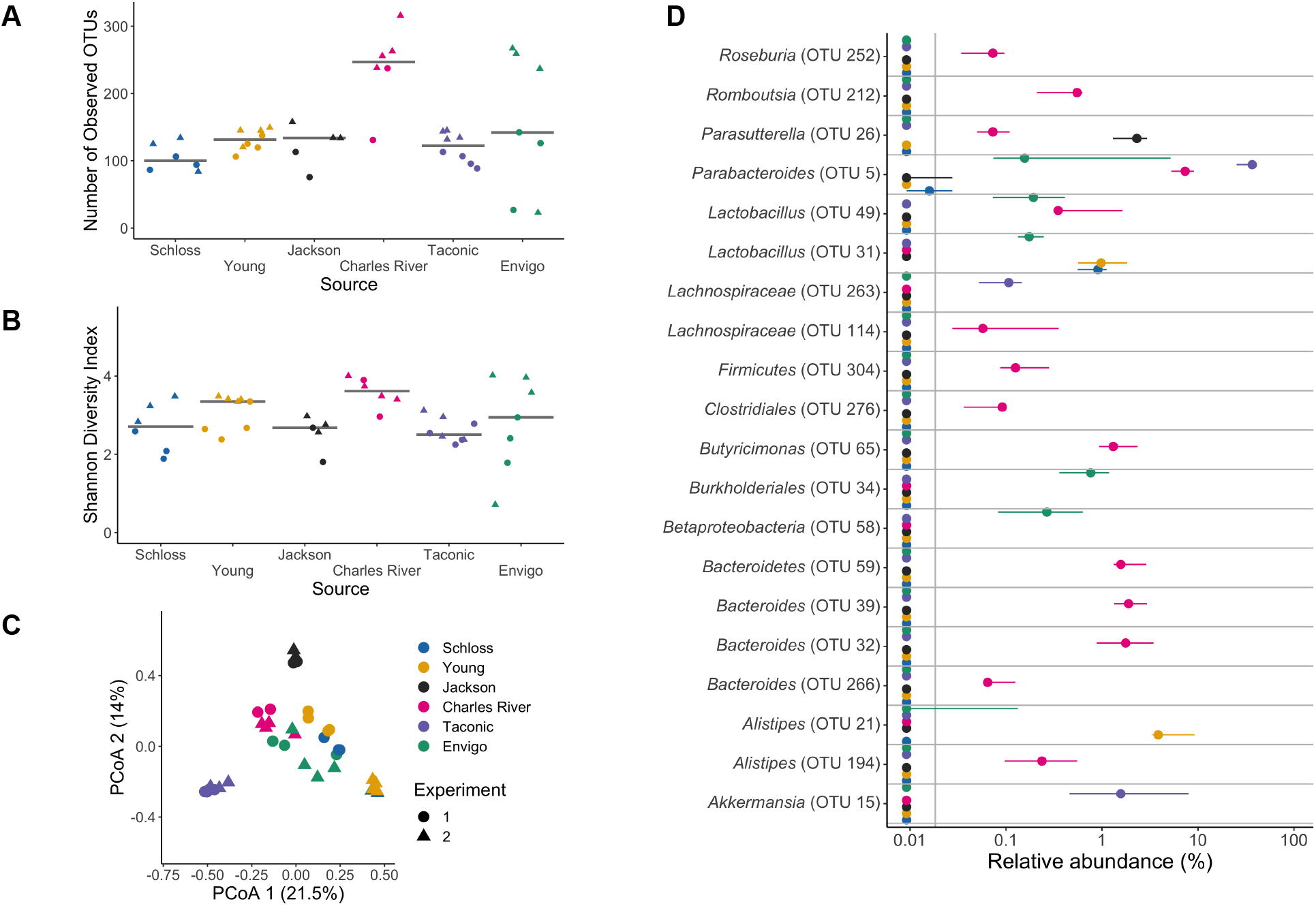
Microbiota variation is high between mice from different sources. A-B. Number of observed OTUs (A) and Shannon diversity index values (B) across sources of mice at baseline (day -1 of the experiment). Differences between sources were analyzed by Kruskal-Wallis test with Benjamini-Hochberg correction for testing each day of the experiment and the adjusted *P* value was < 0.05 for panel A (Table S1). None of the *P* values from pairwise Wilcoxon comparisons between sources were significant after Benjamini-Hochberg correction (Table S2). Gray lines represent the median values for each source of mice. C. Principal Coordinates Analysis (PCoA) of *θ*_*Y C*_ distances of baseline stool samples. Source and the interaction between source and cage effects explained most of the variation (PERMANOVA combined R^2^ = 0.90, *P* < 0.001; Table S3). For A-C: each symbol represents the value for a stool sample from an individual mouse, circles represent experiment 1 mice and triangles represent experiment 2 mice. D. The median (point) and interquantile range (colored lines) of the relative abundances for the 20 most significant OTUs out of the 268 OTUs that varied across sources at baseline (Table S5).

### Clindamycin treatment renders all mice susceptible to *C. difficile* 630 colonization, but clearance time varies across sources

Clindamycin is frequently implicated with human CDIs (42) and was part of the antibiotic treatment for the frequently cited 2008 CDI mouse model (14). We have previously demonstrated mice are rendered susceptible to *C. difficile*, but clear the pathogen within 9 days when treated with clindamycin alone (21, 43). All mice were treated with 10 mg/kg clindamycin via intraperitoneal injection and one day later challenged with 10^3^ *C. difficile* 630 spores (Fig. 2A). The day after infection, *C. difficile* was detectable in all mice at a similar level (median CFU range: 2.2e+07-1.3e+08; *P*_FDR_ = 0.15), indicating clindamycin rendered all mice susceptible regardless of source (Fig. 2B). However, between 3 and 7 days post-infection, we observed variation in *C. difficile* levels across sources of mice (all *P*_FDR_ ≤ 0.019; Fig. 2B and Table S6). This suggested the source of mice was associated with *C. difficile* clearance. While the colonization dynamics were similar between the two experiments, the Schloss mice took longer to clear in the first experiment compared to the second and the Envigo mice took longer to clear in the second experiment compared to the first (Fig. S2A-B). The change in the mice’s weight significantly varied across sources of mice with the most weight loss occurring two days post-infection (Fig. 2C and Table S7). There was also one Jackson and one Envigo mouse that died between 1- and 3-days post-infection during the second experiment. Mice obtained from Jackson, Taconic, and Envigo tended to lose more weight, have higher *C. difficile* CFU levels and take longer to clear the infection compared to the other sources of mice (although there was variation between experiments with Schloss and Envigo mice). This was particularly evident 7 days post-infection (Fig. 2B-C, Fig. S2C-D), when 57% of the mice were still colonized with *C. difficile* (Fig. S2E). By 9 days post-infection the majority of the mice from all sources had cleared *C. difficile* with the exception of 1 Taconic mouse from the first experiment and 2 Envigo mice from the second experiment (Fig. 2B). Thus, clindamycin rendered all mice susceptible to *C. difficile* 630 colonization, regardless of source, but there was significant variation in disease phenotype across the sources of mice.

**Figure 2.**
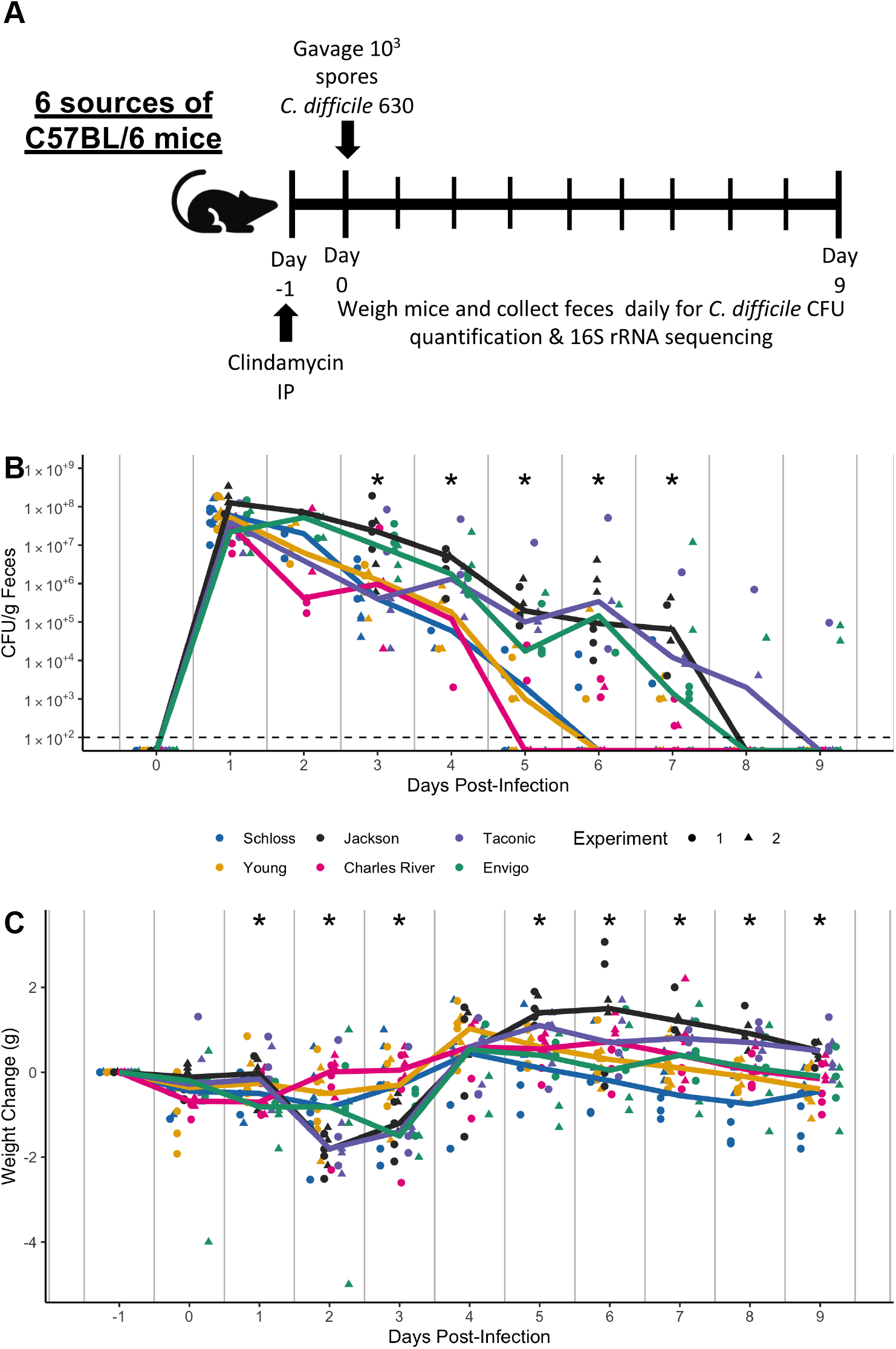
Clindamycin is sufficient to promote *C. difficile* colonization in all mice, but clearance time varies across sources. A. Setup of the experimental timeline. Mice for the experiments were obtained from 6 different sources: the Schloss (N = 8) and Young lab (N = 9) colonies at the University of Michigan, the Jackson Laboratory (N = 8), Charles River Laboratory (N = 8), Taconic Biosciences (N = 8), and Envigo (N = 8). All mice were administered 10 mg/kg clindamycin intraperitoneally (IP) 1 day before challenge with *C. difficile* 630 spores on day 0. Mice were weighed and feces was collected daily through the end of the experiment (9 days post-infection). Note: 3 mice died during course of experiment. 1 Taconic mouse prior to infection and 1 Jackson and 1 Envigo mouse between 1- and 3-days post-infection. B. *C. difficile* CFU/gram stool measured over time (N = 20-49 mice per timepoint) via serial dilutions. The black line represents the limit of detection for the first serial dilution. CFU quantification data was not available for each mouse due to early deaths, stool sampling difficulties, and not plating all of the serial dilutions. C. Mouse weight change measured in grams over time (N = 45-49 mice per timepoint), all mice were normalized to the weight recorded 1 day before infection. For B-C: timepoints where differences between sources of mice were statistically significant by Kruskal-Wallis test with Benjamini-Hochberg correction for testing across multiple days (Table S6 and Table S7) are reflected by the asterisk above each timepoint (*, *P* < 0.05). Lines represent the median for each source and circles represent individual mice from experiment 1 while triangles represent mice from experiment 2.

### Clindamycin treatment alters bacteria in all sources, but a subset of bacterial differences across sources persists

Given the variation in fecal communities that we observed across breeding colonies, we hypothesized that variation in *C. difficile* clearance would be explained by community variation across the 6 sources of mice. As expected, clindamycin treatment decreased the richness and Shannon diversity across all sources of mice (Fig. 3A-B). Interestingly, significant differences in diversity metrics between sources emerged after clindamycin treatment, with Charles River mice having higher richness and Shannon diversity than most of the other sources (*P*_FDR_ < 0.05; Fig 3A-B and Tables S1-2). The clindamycin treatment decreased the variation in community structures between sources of mice. The source of mice and the interactions between source and cage effects explained almost all of the observed variation between communities (combined R^2^ = 0.99, *P* < 0.001; Fig. 3C and Table S3). However, there were only 18 OTUs with relative abundances that significantly varied between sources (Fig. 3D and Table S8). Next, we identified the bacteria that shifted after clindamycin treatment, regardless of source by analyzing paired fecal samples from mice that were collected at baseline and after clindamycin treatment. We identified 153 OTUs that were altered after clindamycin treatment in most mice (Fig. 3E and Table S9). When we compared the list of significant clindamycin impacted bacteria with the bacteria that varied between sources post-clindamycin, we found 4 OTUs that were shared between the lists (*Enterobacteriaceae (OTU 1), Lachnospiraceae* (OTU 130), *Lactobacillus* (OTU 6), *Enterococcus* (OTU 23); Fig. 3D-E and Tables S8-9). Importantly, some of the OTUs that varied between sources also shifted with clindamycin treatment. For example, *Proteus* increased after clindamycin treatment (Fig. 3D), but only in Taconic mice. *Enterococcus* was primarily found only in mice purchased from commercial vendors and also increased in relative abundance after clindamycin treatment (Fig. 3D). These findings demonstrate that clindamycin had a consistent impact on the fecal bacterial communities of mice from all sources and only a subset of the OTUs continued to vary between sources.

**Figure 3.**
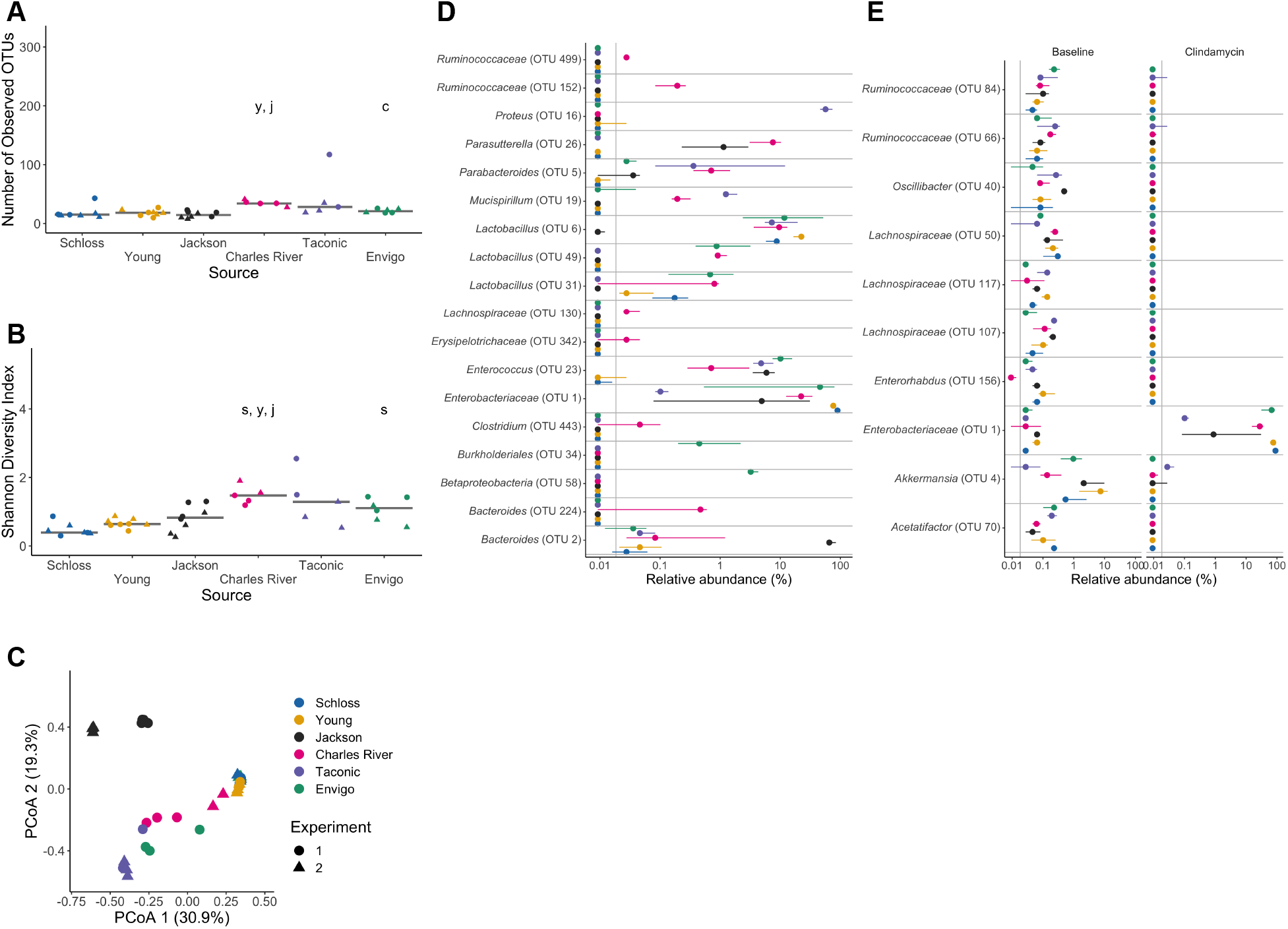
Clindamycin treatment alters bacteria in all sources, but a subset of bacterial differences across sources persists. A-B. Number of observed OTUs (A) and Shannon diversity index values (B) across sources of mice after clindamycin treatment (day 0). Differences between sources were analyzed by Kruskal-Wallis test with Benjamini-Hochberg correction for testing each day of the experiment and the adjusted *P* value was < 0.05 (Table S1). Significant *P* values from the pairwise Wilcoxon comparisons between sources with Benjamini-Hochberg correction are displayed as the first initial of each group compared to the group that they are listed above (Table S2). C. PCoA of *θ*_*Y C*_ distances from stools collected post-clindamycin. Source and the interaction between source and cage effects explained most of the variation observed post-clindamycin (PERMANOVA combined R^2^ = 0.99, *P* < 0.001; Table S3). For A-C, each symbol represents a stool sample from an individual mouse, with circles representing experiment 1 mice and triangles representing experiment 2 mice. D. The median (point) and interquantile range (colored lines) of the relative abundances for the 18 OTUs (Table S8) that varied between sources after clindamycin treatment (day 0). E. The median (point) and interquantile range (colored lines) of the top 10 most significant OTUs out of 153 with relative abundances that changed because of the clindamycin treatment (adjusted *P* value < 0.05). Data were analyzed by paired Wilcoxon signed rank test of mice that had paired sequence data for baseline (day -1) and post-clindamycin (day 0) timepoints (N = 31), with Benjamini-Hochberg correction for testing all identified OTUs (Table S9). The gray vertical line indicates the limit of detection.

### Microbiota variation between sources is maintained after *C. difficile* challenge

One day post-infection, significant differences in diversity metrics remained across sources (*P*_FDR_ < 0.05, Fig 4A-B and Tables S1-2). Although the Charles River mice had more diverse communities and were also able to clear *C. difficile* faster than the other sources, diversity did not explain the observed variation in *C. difficile* colonization across sources. The Young and Schloss mice had the lowest diversity 1 day post-infection and were able to clear *C. difficile* earlier than Jackson, Taconic and Envigo mice. The source of mice and the interactions between source and cage effects continued to explain most of the observed community variation (combined R^2^ = 0.88; *P* < 0.001; Fig. 4C and Table S3). One day after *C. difficile* challenge, there were 44 OTUs with significantly different relative abundances across sources (Fig. 4D and Table S10).

**Figure 4.**
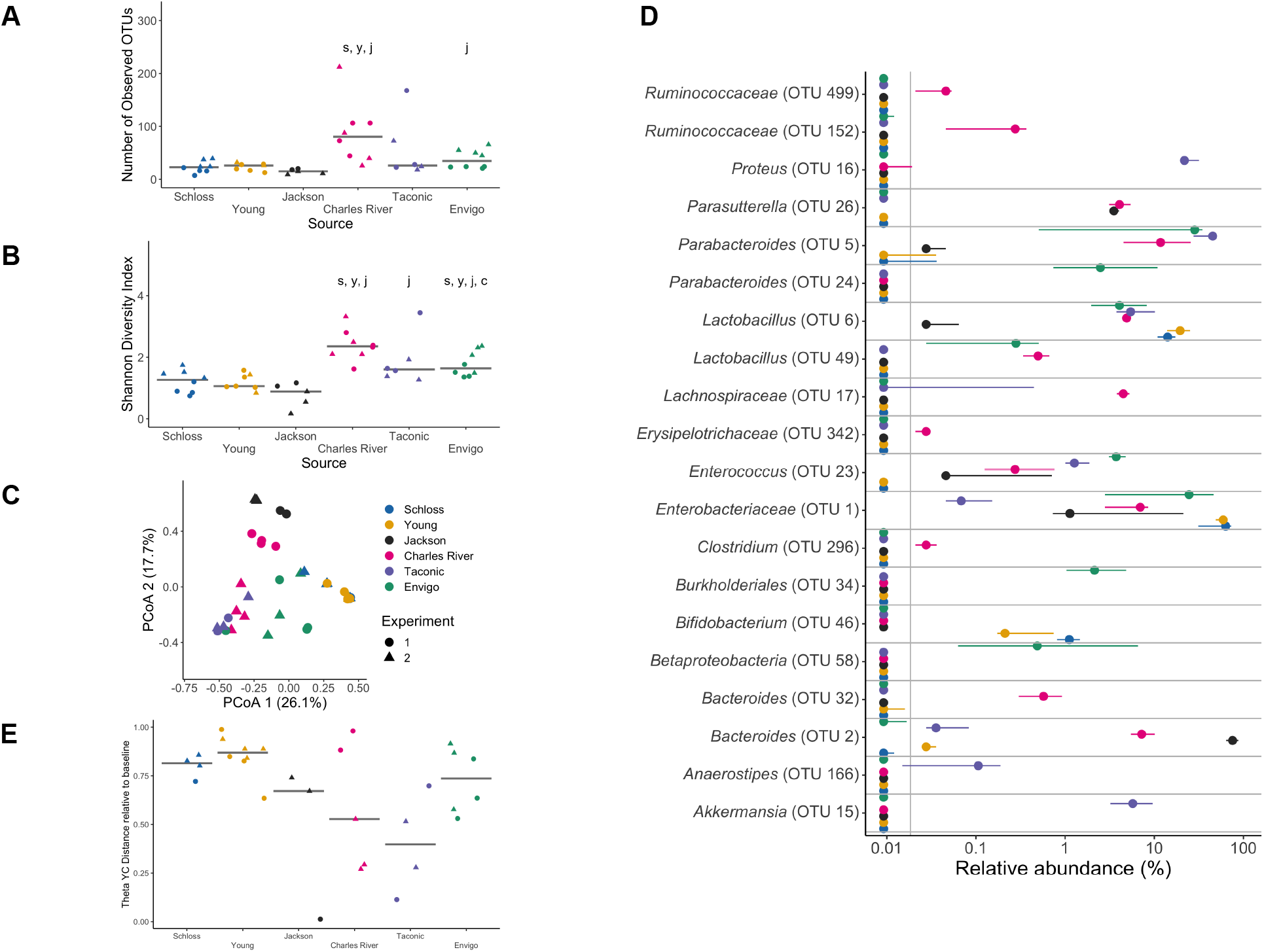
Microbiota variation across sources is maintained after *C. difficile* challenge. A-B. Number of observed OTUs (A) and Shannon diversity index values (B) across sources of mice 1-day post-infection. Data were analyzed by Kruskal-Wallis test with Benjamini-Hochberg correction for testing each day of the experiment and the adjusted *P* value was < 0.05 (Table S1). Significant *P* values from the pairwise Wilcoxon comparisons between sources with Benjamini-Hochberg correction are displayed as the first initial of each group compared to the group that they are listed above (Table S2). PCoA of *θ*_*Y C*_ distances of 1-day post-infection stool samples. Source and the interaction between source and cage effects explained most of the variation between fecal communities (PERMANOVA combined R^2^ = 0.88, *P* < 0.001; Table S3). For A-C: each symbol represents the value for a stool sample from an individual mouse, circles represent experiment 1 mice and triangles represent experiment 2 mice. D. The median (point) and interquantile range (colored lines) of the relative abundances for the top 20 most significant OTUs out of the 44 OTUs that varied between sources 1-day post-infection. The gray vertical line indicates the limit of detection. For each timepoint OTUs with differential relative abundances across sources of mice were identified by Kruskal-Wallis test with Benjamini-Hochberg correction for testing all identified OTUs (Table S10). E. *θ*_*Y C*_ distances of fecal samples collected 7-days post-infection relative to the baseline (day -1) sample for each mouse. Each symbol represents an individual mouse. Gray lines represent the median for each source.

Throughout the experiment, the source of mice continued to be the dominant factor that explained the observed variation across fecal communities (PERMANOVA R^2^ = 0.35, *P* < 0.001) followed by interactions between cage effects and the day of the experiment (Movie S1 and Table S11). Fecal samples from the same source of mice continued to cluster closely to each other throughout the experiment. By 7 days post-infection, when approximately 43% mice had cleared *C. difficile*, most of the mice had not recovered to their baseline community structure (Fig. 4E). The distance to the baseline community did not explain the variation in *C. difficile* clearance as the Schloss and Young mice had mostly cleared *C. difficile*, but their communities were a greater distance from baseline 7 days post-infection compared to the Jackson and Taconic mice that were still colonized. In summary, mouse bacterial communities varied significantly between sources throughout the course of the experiment and a consistent subset of bacteria remained different between sources regardless of clindamycin and *C. difficile* challenge.

### Baseline, post-clindamycin, and post-infection community data can predict mice that will clear *C. difficile* by 7 days post-infection

After identifying taxa that varied between sources, changed after clindamycin treatment, or both, we determined which taxa were influencing the variation in *C. difficile* colonization at day 7 (Fig. 2B, Fig. S2C). We trained three L2-regularized logistic regression models with either input bacterial community data from the baseline (day = -1), post-clindamycin (day = 0), or post-infection (day = 1) timepoints of the experiment to predict *C. difficile* colonization status on day 7 (Fig. S3A-B). All models were better at predicting *C. difficile* colonization status on day 7 than random chance (all *P* < 0.001, Table S12). The model based on the post-clindamycin (AUROC = 0.78) community OTU data performed significantly better than the baseline (AUROC = 0.72) or the post-infection (AUROC = 0.67) models (*P*_FDR_ < 0.001 for pairwise comparisons; Fig. S3C and Table S13). Thus, we were able to use bacterial relative abundance data from the time of *C. difficile* challenge to differentiate mice that had cleared *C. difficile* before day 7 from the mice still colonized with *C. difficile* at that timepoint. This result suggests that the bacterial community’s response to clindamycin treatment had the greatest influence on subsequent *C. difficile* colonization dynamics.

To examine the bacteria that were driving each model’s performance, we selected the 20 OTUs that had the highest absolute feature weights in each of the 3 models (Table S14). First, we looked at OTUs from the model with the best performance, which was based on the post-clindamycin treatment (day 0) bacterial community data. Out of the 10 highest ranked OTUs, 7 OTUs were associated with *C. difficile* colonization 7 days post-infection (*Bacteroides, Escherichia/Shigella*, 2 *Lachnospiraceae, Lactobacillus, Porphyromonadaceae*, and *Ruminococcaceae*), while 3 OTUs were associated with clearance (*Enterobacteriaceae, Lachnospiraceae, Porphyromonadaceae*; Fig. 5A). Next, we examined whether any of the top 20 ranked OTUs from the post-clindamycin (day 0) model were also important in the other 2 classification models based on baseline (day -1) and 1 day post-infection community data. We identified 6 OTUs that were important to the post-clindamycin model and either the baseline or 1 day post-infection models (*Enterobacteriaceae, Ruminococcaceae, Lactobacillus, Bacteroides, Porphyromonadaceae, Erysipelotrichaceae*; Table S14). Thus, a subset of bacterial OTUs were important for determining *C. difficile* colonization dynamics across multiple timepoints.

**Figure 5.**
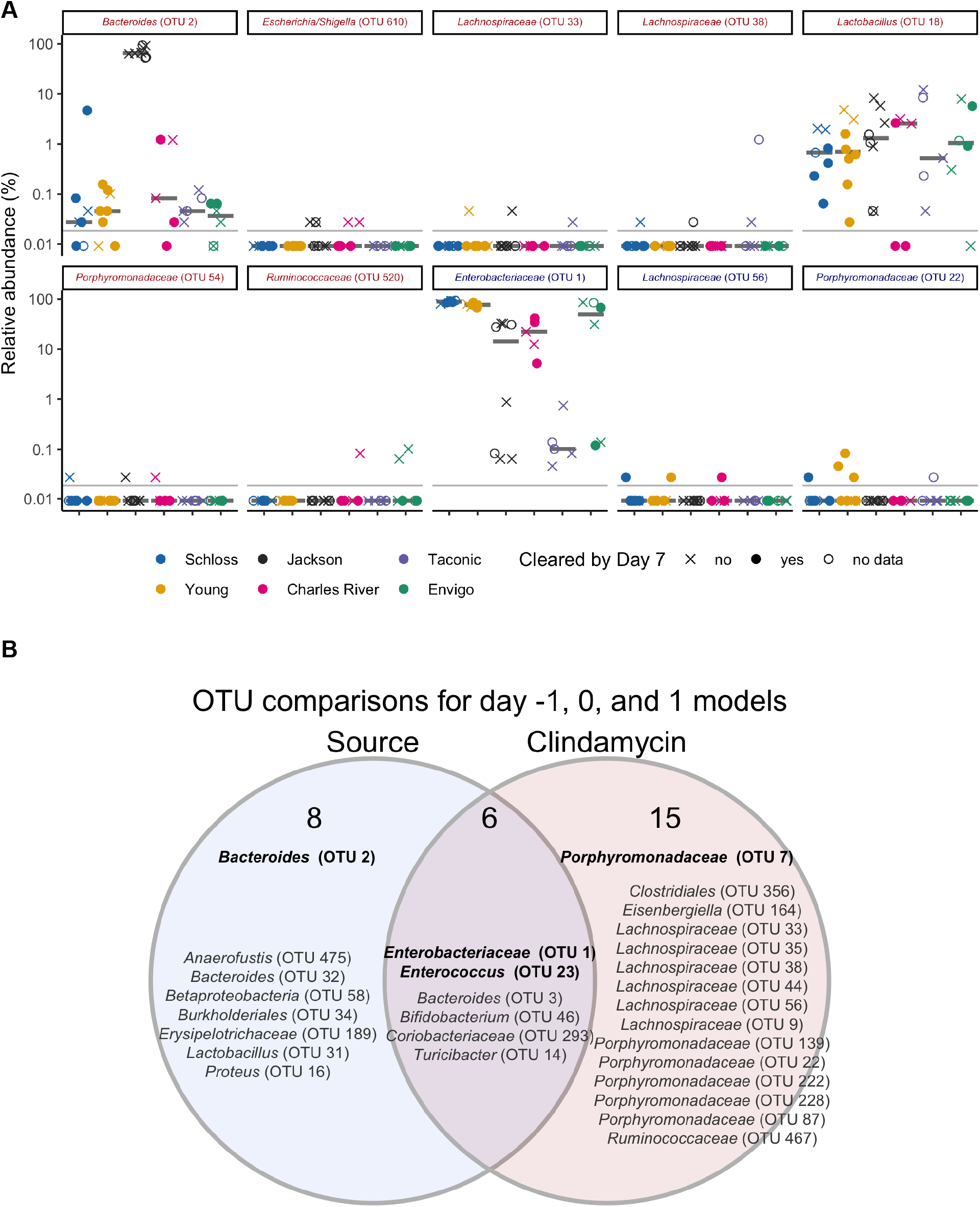
Bacteria that influenced whether mice cleared *C. difficile* by day 7. A. Post-clindamycin (day 0) relative abundance data for the 10 OTUs with the highest rankings based on feature weights in the post-clindamycin (day 0) classification model. Red font represents OTUs that correlated with *C. difficile* colonization and blue font represents OTUs that correlated with clearance. Symbols represent the relative abundance data for an individual mouse. Gray bars indicate the median relative abundances for each source. B. Venn diagram that combines OTUs that were important to the day -1, 0, and 1 classification models (Fig. S4, Table S14) and either overlapped with taxa that varied across sources at the same timepoint, were impacted by clindamycin treatment, or both. Bold OTUs were important to more than 1 classification model.

To determine whether the OTUs driving the classification models also varied between sources, were altered by clindamycin treatment, or both, we identified the OTUs from each model that varied between sources (Fig. 1D, 3D, 4D and Tables S5, S8, S10) or were impacted by clindamycin treatment (Fig. 3E and Table S9; Fig. S4). Comparing the features important to the 3 models identified 14 OTUs associated with source, 21 OTUs associated with clindamycin treatment, and 6 OTUs associated with both (Fig. 5B). Together, these results suggest that the initial bacterial communities and their responses to clindamycin influenced the clearance of *C. difficile*.

Several OTUs that overlapped with our previous analyses appeared across at least 2 models (*Bacteroides, Enterococcus, Enterobacteriaceae, Porphyromonadaceae*), so we examined how the relative abundances of these OTUs varied over the course of the experiment (Fig. 6). Across the 9 days post-infection, there was at least 1 timepoint when the relative abundances of these OTUs significantly varied between sources (Table S15). Interestingly, there were no OTUs that emerged as consistently enriched or depleted in mice that were colonized past 7 days post-infection, suggesting that multiple bacteria influence *C. difficile* colonization dynamics.

**Figure 6:**
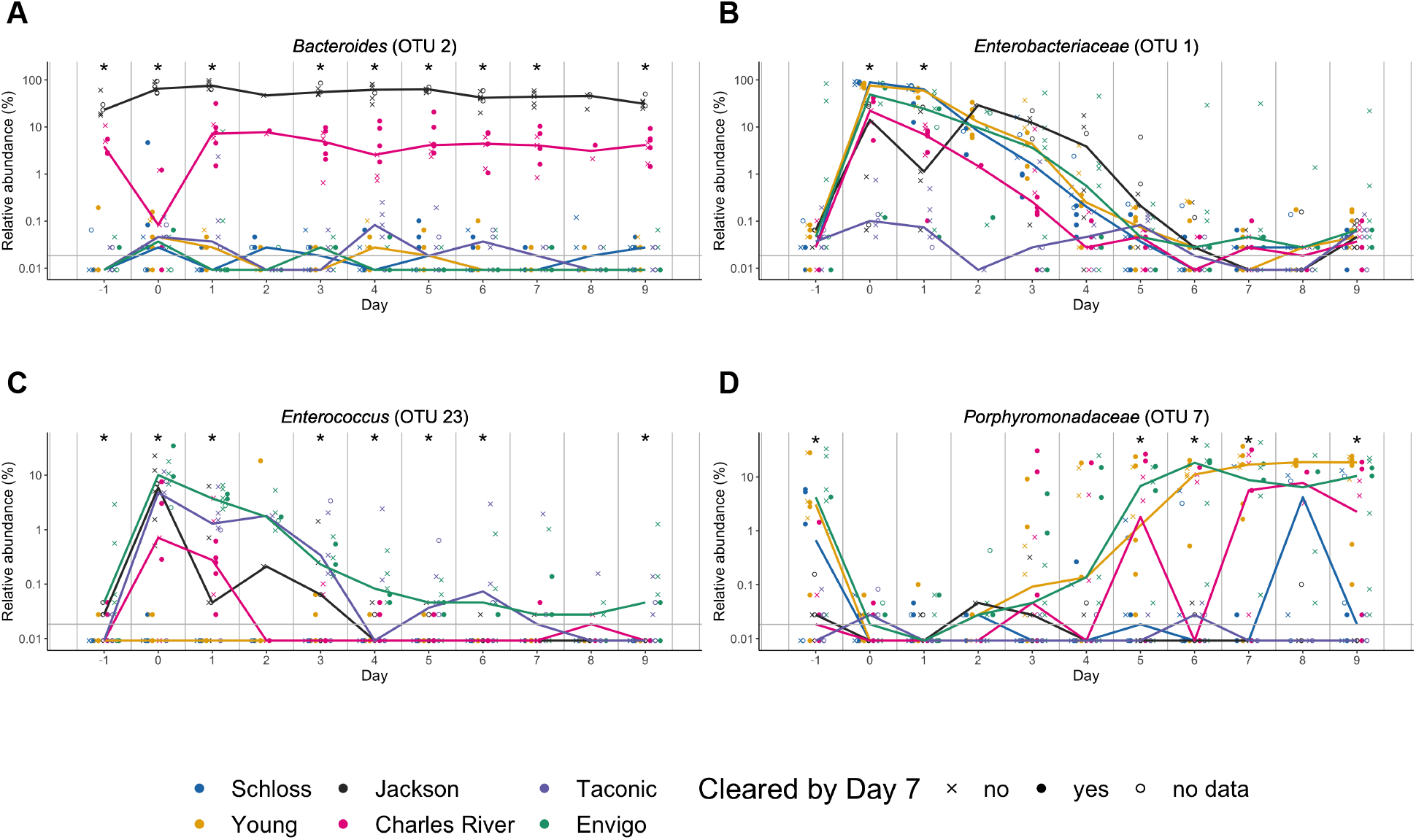
OTUs associated with *C. difficile* colonization dynamics vary across sources throughout the experiment. A-D. Relative abundances of bold OTUs from Fig. 5B that were important in at least two classification models are shown over time. A. *Bacteroides* (OTU 2), which varied across sources throughout the experiment. B-C. *Enterobacteriaceae* (B) and *Enterococcus* (C), which significantly varied across sources and were impacted by clindamycin treatment. D. *Porphyromonadaceae* (OTU 7), which was significantly impacted by clindamycin treatment and after examining relative abundance dynamics over the course of the experiment was found to also significantly vary between sources of mice on days -1, 5, 6, 7, and 9 of the experiment. Symbols represent the relative abundance data for an individual mouse. Colored lines indicate the median relative abundances for each source. The gray horizontal line represents the limit of detection. Timepoints where differences between sources of mice were statistically significant by Kruskal-Wallis test with Benjamini-Hochberg correction for testing across multiple days (Table S15) are identified by the asterisk above each timepoint (*, P < 0.05).

## Discussion

Applying our CDI model to 6 different sources of mice, allowed us to identify bacterial taxa that were unique to different sources as well as taxa that were universally impacted by clindamycin. We trained logistic regression models with baseline (day -1), post-clindamycin treatment (day 0), and 1-day post-infection fecal community data that could predict whether mice cleared *C. difficile* by 7 days post-infection better than random chance. We identified *Bacteroides, Enterococcus, Enterobacteriaceae, Porphyromonadaceae* (Fig. 6) as candidate bacteria within these communities that influenced variation in *C. difficile* colonization dynamics since these bacteria were all important in the logistic regression models and varied by source, were impacted by clindamycin treatment, or both. Overall, our results demonstrated clindamycin was sufficient to render mice from multiple sources susceptible to CDI and only a subset of the inter-individual microbiota variation across mice from different sources was needed to predict which mice could clear *C. difficile*.

Other studies have used mice from multiple sources to identify bacteria that either promote colonization resistance or increase susceptibility to enteric infections (22, 23, 26–30). For example, against *Salmonella* infections, *Enterobacteriaceae* and segmented filamentous bacteria have emerged as protective (22, 27). We found *Enterobacteriaceae* increased in all sources of mice after clindamycin treatment, facilitating *C. difficile* colonization. However, there was also variation in *Enterobacteriaceae* relative abundance levels between sources that was associated with the variation in *C. difficile* colonization dynamics across sources. Thus, bacteria may have differential roles in determining susceptibility depending on the type of bacterial infection.

Differences in CDI mouse model studies have been attributed to intestinal microbiota variation across sources. For example, researchers using the same clindamycin treatment and C57BL/6 mice had different *C. difficile* outcomes, one having sustained colonization (32), while the other had transient colonization (18), despite both using *C. difficile* VPI 10643. Baseline differences in the microbiota composition have been hypothesized to partially explain the differences in colonization outcomes and overall susceptibility to *C. difficile* after treatment with the same antibiotic (13, 31). When we treated mice from 6 different sources with clindamycin and challenged them with *C. difficile* 630, we found microbiota variation across sources impacted colonization outcomes, but not susceptibility. A previous study with *C. difficile* identified an endogenous protective *C. difficile* strain LEM1 that bloomed after antibiotic treatment in mice from Jackson or Charles River Laboratories, but not Taconic that protected mice against the more toxigenic *C. difficile* VPI10463 (26). Given that we obtained mice from the same vendors, we checked all mice for endogenous *C. difficile* by plating stool samples that were collected after clindamycin treatment. However, we did not identify any endogenous *C. difficile* strains prior to challenge, suggesting there were no endogenous protective strains in the mice we received and other bacteria mediated the variation in *C. difficile* colonization across sources. The *C. difficile* strain used could also be contributing to the variation in *C. difficile* outcomes seen across different research groups. For example, a group found differential colonization outcomes after clindamycin treatment, with *C. difficile* 630 and M68 infections eventually becoming undetectable while strain BI-7 remained detectable up to 70 days post-treatment (44). The bacterial perturbations induced by clindamycin treatment have been well characterized and our findings agree with previous CDI mouse model work demonstrating *Enterococcus* and *Enterobacteriaceae* were associated with *C. difficile* susceptibility and *Porpyhromonadaceae, Lachnospiraceae, Ruminococcaceae*, and *Turicibacter* were associated with resistance (19, 21, 32, 33, 43–46). While we have demonstrated that susceptibility is uniform across sources of mice after clindamycin treatment, there could be different outcomes for either susceptibility or clearance in the case of other antibiotic treatments.

We found the time needed to naturally clear *C. difficile* varied across sources of mice implying that at least in the context of the same perturbation, microbiota differences influence infection outcome. More importantly, we were able to explain the variation observed across sources with a subset of OTUs that were also important for predicting *C. difficile* colonization status 7 days post-infection. Since all but 3 mice eventually cleared *C. difficile* 630 by 9 days post-infection and the model built with the post-clindamycin (day 0) OTU relative abundance data had the best performance, our results suggest clindamycin treatment had a larger role in determining *C. difficile* susceptibility and clearance than the source of the mice.

Using mice from multiple sources successfully increased the inter-animal variation. One alternative approach that has been used in some CDI studies is to associate mice with human microbiotas (47–52). However, a major caveat to this method is the substantial loss of human microbiota community members upon transfer to mice (53, 54). Additionally with the exception of 2 recent studies (47, 48), most of these studies associated mice with just 1 types of human microbiota either from a single donor or a single pool from multiple donors (49–52). This approach does not aid in the goal of modeling the interpersonal variation seen in humans to understand how the microbiota influences susceptibility to CDIs and adverse outcomes. Importantly, our study using mice from 6 different sources increased the variation between groups of mice compared to using 1 source alone, to better reflect the inter-individual microbiota variation observed in humans.

Another motivation for associating mice with human microbiotas is to study the bacteria associated with the disease in humans. Decreased *Bifidobacterium, Porphyromonas, Ruminococcaceae* and *Lachnospiraceae* and increased *Enterobacteriaceae, Enterococcus, Lactobacillus*, and *Proteus* have all been associated with human CDIs (7). Encouragingly, these populations were well represented in our study, suggesting most of the mouse sources are suitable for gaining insights into the bacteria influencing *C. difficile* colonization and infections in humans. An important exception was *Enterococcus*, which was primarily absent from University of Michigan colonies and *Proteus*, which was only found in Taconic mice. The fact that some CDI-associated bacteria were only found in a subset of mice has important implications for future CDI mouse model studies, but also models the natural patchiness of microbial populations in humans.

Other microbiota and host factors that were outside the scope of our current study may also contribute to the differences in *C. difficile* colonization dynamics between sources of mice. The microbiota is composed of viruses, fungi, and parasites in addition to bacteria, and these non-bacterial members can also vary across sources of mice (55, 56). While our study focused solely on the bacterial portion, viruses and fungi have also begun to be implicated in the context of CDIs or FMT treatments for recurrent CDIs (35, 57–60). Beyond community composition, the metabolic function of the microbiota also has a CDI signature (20, 46, 61, 62) and can vary across mice from different sources (63). For example, microbial metabolites, particularly secondary bile acids and butyrate production, have been implicated as important contributors to *C. difficile* resistance (33, 44). Interestingly, butyrate has previously been shown to vary across mouse vendors and mediated resistance to *Citrobacter rodentium* infection, a model of enterohemorrhagic and enteropathogenic *Escherichia coli* infections (23). Evidence for immunological toning differences in IgA and Th17 cells across mice from different vendors have also been documented and (64, 65) could influence the host response to CDI (66, 67). The outcome after *C. difficile* exposure depends on a multitude of factors, including age, diet, and immunity; all of which are also influenced by the microbiota.

We have demonstrated that the ways baseline microbiotas from different mouse sources respond to clindamycin treatment influence the length of time mice remained colonized with *C. difficile* 630. To better understand the contribution of the microbiota to *C. difficile* pathogenesis and treatments, using multiple sources of mice may yield more insights than a single source. Furthermore, for studies wanting to examine the interplay between particular bacteria such as *Enterococcus* and *C. difficile*, these results could serve as a resource for selecting mice to address the question. Using mice from multiple sources helps model the interpersonal microbiota variation among humans to aid our understanding of how the gut microbiota provides colonization resistance to CDIs.

## Acknowledgements

This work was supported by the National Institutes of Health (U01AI124255). ST was supported by the Michigan Institute for Clincial and Health Research Postdoctoral Translation Scholars Program (UL1TR002240 from the National Center for Advancing Translational Sciences). We thank members of the Schloss lab for feedback on planning the experiments and data presentation. In particular, we thank Begüm Topçuoğlu for help with implementing logistic regression models, Ana Taylor for help with media preparation and sample collection, and Nicholas Lesniak for his critical feedback on the manuscript. We also thank members of Vincent Young’s lab, particularly Kimberly Vendrov, for guidance with the *C. difficile* infection mouse model and donating the mice. We also thank the Unit for Laboratory Animal Medicine at the University of Michigan for maintaining our mouse colony and providing the institutional support for our mouse experiments. Finally, we thank Kwi Kim, Austin Campbell, and Kimberly Vendrov for their help in maintaining the Schloss lab’s anaerobic chamber.

## Materials and Methods

### (i) Animals

All experiments were approved by the University of Michigan Animal Care and Use Committee (IACUC) under protocol number PRO00006983. Female C57BL/7 mice were obtained from 6 different sources: The Jackson Laboratory, Charles River Laboratories, Taconic Biosciences, Envigo, and two colonies at the University of Michigan (the Schloss lab colony and the Young lab colony). The Young lab colony was originally established with mice purchased from Jackson in 2002, and the Schloss lab colony was established in 2010 with mice donated from the Young lab. The 4 groups of mice purchased from vendors were allowed to acclimate to the University of Michigan mouse facility for 13 days prior to starting the experiment. At least 4 female mice (age 5-10 weeks) were obtained per source and mice from the same source were primarily housed at a density of 2 mice per cage. The experiment was repeated once, approximately 3 months after the start of the first experiment.

### (ii) Antibiotic treatment

After the 13-day acclimation period, all mice received 10 mg/kg clindamycin (filter sterilized through a 0.22 micron syringe filter prior to administration) via intraperitoneal injection (Fig. 1A).

### (iii) *C. difficile* infection model

Mice were challenged with 10^3^ spores of *C. difficile* strain 630 via oral gavage post-infection 1 day after clindamycin treatment as described previously (21). Mice weights and stool samples were taken daily through 9 days post-challenge (Fig. 1A). Collected stool was split for *C. difficile* quantification and 16S rRNA sequencing analysis. For *C. difficile* quantification, stool samples were transferred to the anaerobic chamber, serially diluted in PBS, plated on taurocholate-cycloserine-cefoxitin-fructose agar (TCCFA) plates, and counted after 24 hours of incubation at 37°C under anaerobic conditions. A sample from the day 0 timepoint (post-clindamycin and prior to *C. difficile* challenge) was also plated on TCCFA to ensure mice were not already colonized with *C. difficile* prior to infection. There were 3 deaths recorded over the course of the experiment, 1 Taconic mouse died prior to *C. difficile* challenge and 1 Jackson and 1 Envigo mouse died between 1- and 3-days post-infection. Mice were categorized as cleared when no *C. difficile* was detected in the first serial dilution (limit of detection: 100 CFU). Stool samples for 16S rRNA sequencing were snap frozen in liquid nitrogen and stored at -80°C until DNA extraction.

### (iv) 16S rRNA sequencing

DNA was extracted from -80°C stored stool samples using the DNeasy Powersoil HTP 96 kit (Qiagen) and an EpMotion 5075 automated pipetting system (Eppendorf). The V4 region was amplified for 16S rRNA with the AccuPrime Pfx DNA polymerase (Thermo Fisher Scientific) using custom barcoded primers, as previously described (68). The ZymoBIOMICS microbial community DNA standards was used as a mock community control (69) and water was used as a negative control per 96-well extraction plate. The PCR amplicons were cleaned up and normalized with the SequalPrep normalization plate kit (Thermo Fisher Scientific). Amplicons were pooled and quantified with the KAPA library quantification kit (KAPA biosystems), prior to sequencing using the MiSeq system (Illumina).

### (v) 16S rRNA gene sequence analysis

mothur (v. 1.43) was used to process all sequences (70) with a previously published protocol (68). Reads were combined and aligned with the SILVA reference database (71). Chimeras were removed with the VSEARCH algorithm and taxonomic assignment was completed with a modified version (v16) of the Ribosomal Database Project reference database (v11.5) (72) with an 80% confidence cutoff. Operational taxonomic units (OTUs) were assigned with a 97% similarity threshold using the opticlust algorithm (73). To account for uneven sequencing across samples, samples were rarefied to 5,437 sequences 1,000 times for alpha and beta diversity analyses, and a single time to generate relative abundances for model training. PCoAs were generated based on *θ*_*Y C*_ distances. Permutational multivariate analysis of variance (PERMANOVA) was performed on mothur-generated *θ*_*Y C*_ distance matrices with the adonis function in the vegan package (74) in R (75).

### (vi) Classification model training and evaluation

Models were generated based on mice that were categorized as either cleared or colonized 7 days post-infection and had sequencing data from the baseline (day -1), post-clindamycin (day 0), and post-infection (day 1) timepoints of the experiment. Input bacterial community relative abundance data at the OTU level from the baseline, post-clindamycin, and 1-day post-infection timepoints was used to generate 3 classification models that predicted *C. difficile* colonization status 7 days post-infection. The L2-regularized logistic regression models were trained and tested using the caret package (76) in R as previously described (77) with the exception that we used 60% training and 40% testing data splits for the cross-validation of the training data to select the best cost hyperparameter and the testing of the held out test data to measure model performance. The modified training to testing ratio was selected to accommodate the small number of samples in the dataset. Code was modified from https://github.com/SchlossLab/ML_pipeline_microbiome to update the classification outcomes and change the data split ratios. The modified repository to regenerate our modeling analysis is available at https://github.com/tomkoset/ML_pipeline_microbiome.

### (vii) Statistical analysis

All statistical tests were performed in R (v 4.0.2) (75). The Kruskal-Wallis test was used to analyze differences in *C. difficile* CFU, mouse weight change, and alpha diversity across sources with a Benjamini-Hochberg correction for testing multiple timepoints, followed by pairwise Wilcoxon comparisons with Benjamini-Hochberg correction. For taxonomic analysis and generation of logistic regression model input data, *C. difficile* (OTU 20) was removed. Bacterial relative abundances that varied across sources at the OTU level were identified with the Kruskal-Wallis test with Benjamini-Hochberg correction for testing all identified OTUs, followed by pairwise Wilcoxon comparisons with Benjamini-Hochberg correction. The Wilcoxon rank sum test was used to test for OTUs that differed between experiments within the Schloss, Young, and Envigo sources with Benjamini-Hochberg correction for testing all identified OTUs. OTUs impacted by clindamycin treatment were identified using the paired Wilcoxon signed rank test with matched pairs of mice samples from day -1 and day 0. To determine whether classification models had better performance (test AUROCs) than random chance (0.5), we used the one-sample Wilcoxon signed rank test. To examine whether there was an overall difference in predictive performance across the 3 classification models we used the Kruskal-Wallis test followed by pairwise Wilcoxan comparisons with Benjamini-Hochberg correction for multiple hypothesis testing. The tidyverse package (v 1.3.0) was used to wrangle and graph data (78).

### (viii) Code availability

Code for all data analysis and generating this manuscript is available at https://github.com/SchlossLab/Tomkovich_Vendor_XXXX_2020.

### (ix) Data availability

The 16S rRNA sequencing data have been deposited in the National Center for Biotechnology Information Sequence Read Archive (BioProject Accession no. PRJNA608529).

## Figures

**Figure S1.**
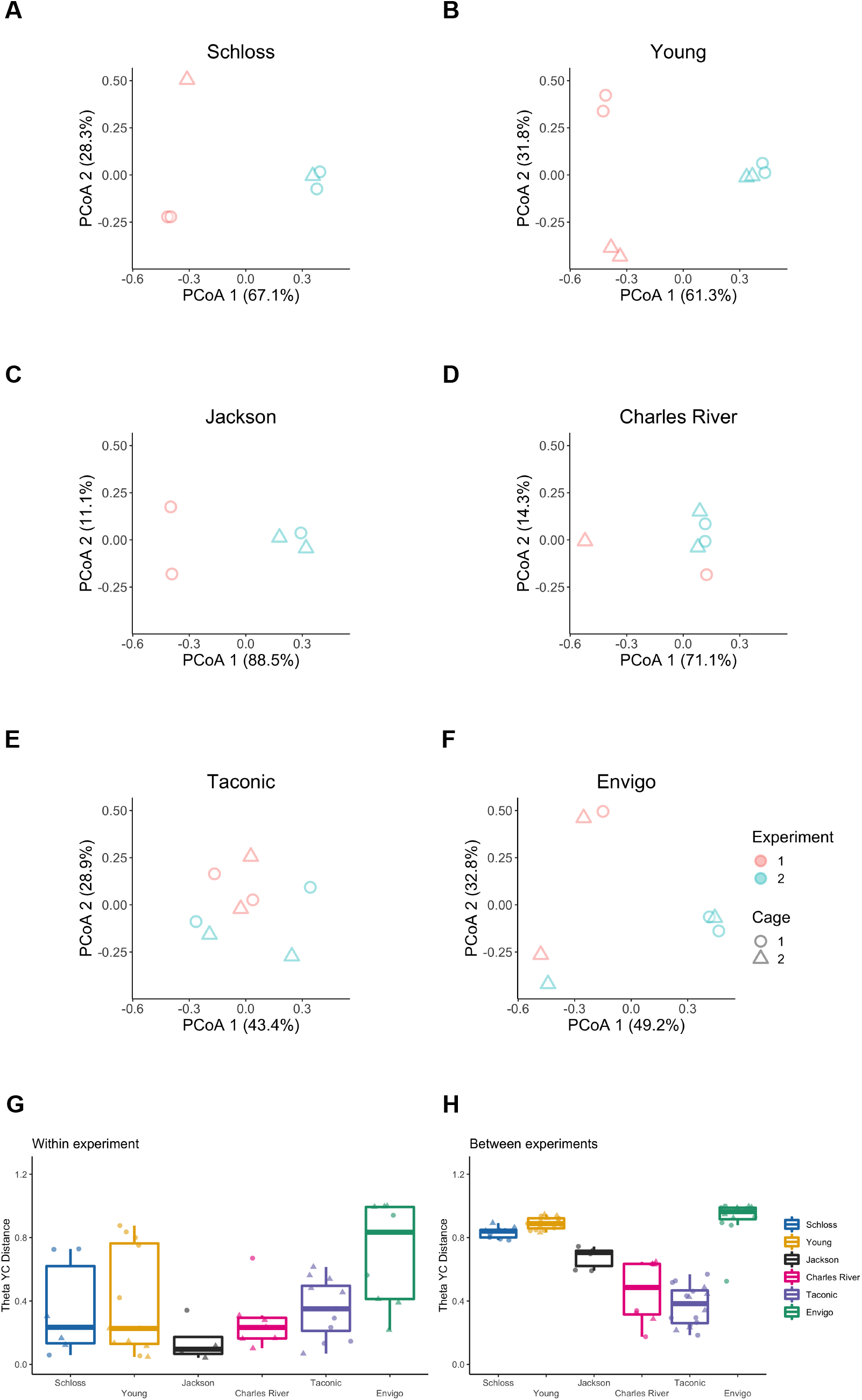
Bacterial communities vary between experiments for some sources. A-F. PCoA of *θ*_*Y C*_ distances for the baseline fecal bacterial communities within each source of mice. Each symbol represents a stool sample from an individual mouse with color corresponding to experiment and shape representing cage mates. Experiment number and cage effects explained most of the observed variation for samples from the Schloss (PERMANOVA combined R^2^ = 0.99; *P ≤* 0.033) and Young (combined R^2^ = 0.95; *P ≤* 0.03) mice (Table S4). G-H: Boxplots of the *θ*_*Y C*_ distances of the 6 sources of mice relative to mice within the same source and experiment (G) or mice within the same source and between experiments (H) at baseline (day -1). Symbols represent individual mouse samples: circles for experiment 1 and triangles for experiment 2.

**Figure S2.**
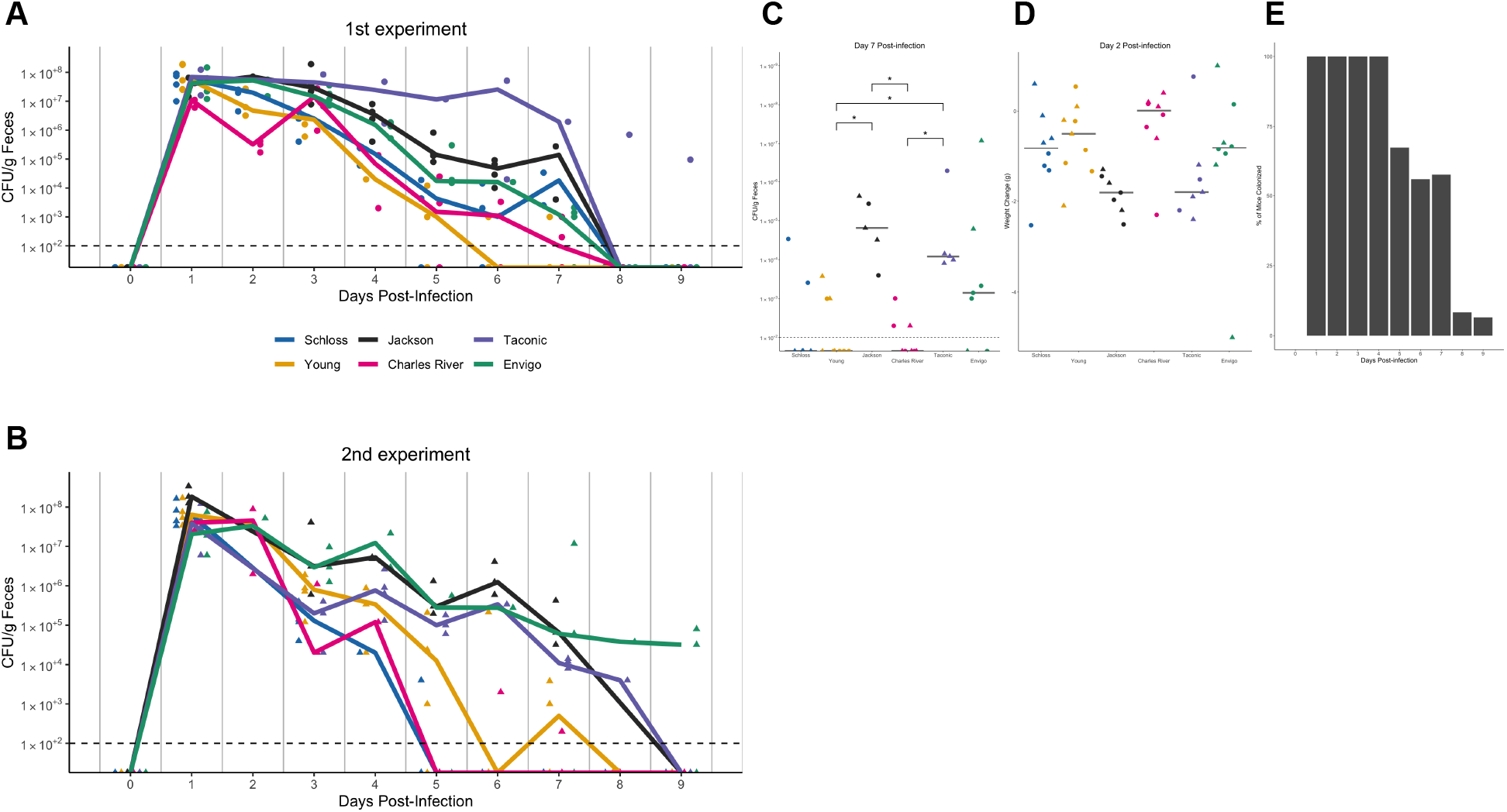
*C. difficile* CFU variation across sources varies slightly between the 2 experiments. A-B. *C. difficile* CFU/gram of stool quantification over time for experiment 1 (A) and 2 (B). Experiments were conducted approximately 3 months apart. Lines represent the median CFU for each source, symbols represent individual mice and the black line represents the limit of detection. C. *C. difficile* CFU/gram stool 7-days post-infection across sources of mice with an asterisk for pairwise Wilcoxon comparisons with Benjamini-Hochberg correction where *P* < 0.05. D. Mouse weight change 2-days post-infection across sources of mice, no pairwise Wilcoxon comparisons were significant after Benjamini-Hochberg correction. For C-D: circles represent experiment 1 mice, triangles represent experiment 2 mice and gray lines indicate the median values for each group. E. Percent of mice that were colonized with *C. difficile* over the course of the experiment. Each day the percent is calculated based on the mice where *C. difficile* CFU was quantified for that particular day. Total N for each day: day 1 (N = 42), day 2 (N = 20), day 3 (N = 39), day 4 (N = 29), day 5 (N = 43), day 6 (N = 34), day 7 (N = 40), day 8 (N = 36), and day 9 (N = 46).

**Figure S3.**
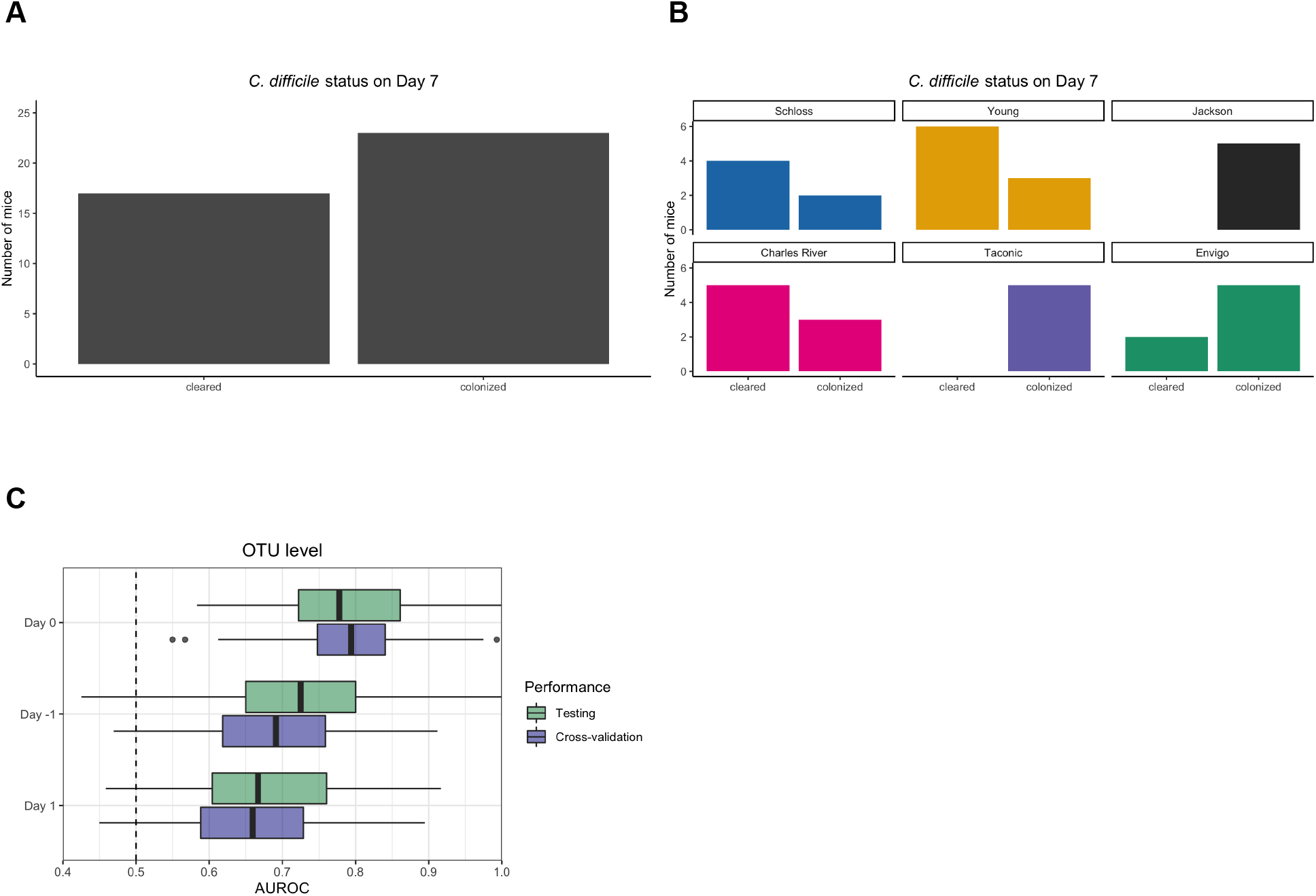
Bacterial community composition before, after clindamycin perturbation, and post-infection can predict *C. difficile* colonization status 7 days post-challenge. A. Bar graph visualizations of overall 7-days post-infection *C. difficile* colonization status that were used as classification outcomes to build L2-regularized logistic regression models. Mice were classified as colonized or cleared (not detectable at the limit of detection of 100 CFU) based on CFU g/stool data from 7 days post-infection. B. *C. difficile* CFU status on Day 7 within each mouse source. N = 8-9 mice per group. C. L2-regularized logistic regression classification model area under the receiving operator characteristic curve (AUROCs) to predict *C. difficile* CFU on day 7 post-infection (Fig. 2B, Fig. S2C) based on the OTU community relative abundances at baseline (day -1), post-clindamycin (day 0), and 1-day post-infection. All models performed better than random chance (AUROC = 0.5, all *P* < 0.001, Table S12) and the model built with post-clindamycin bacterial OTU relative abundances had the best performance ((*P*_FDR_ < 0.001 for all pairwise comparisons, Table S13). See Table S14 for list of the 20 OTUs that were ranked as most important to each model.

**Figure S4.**
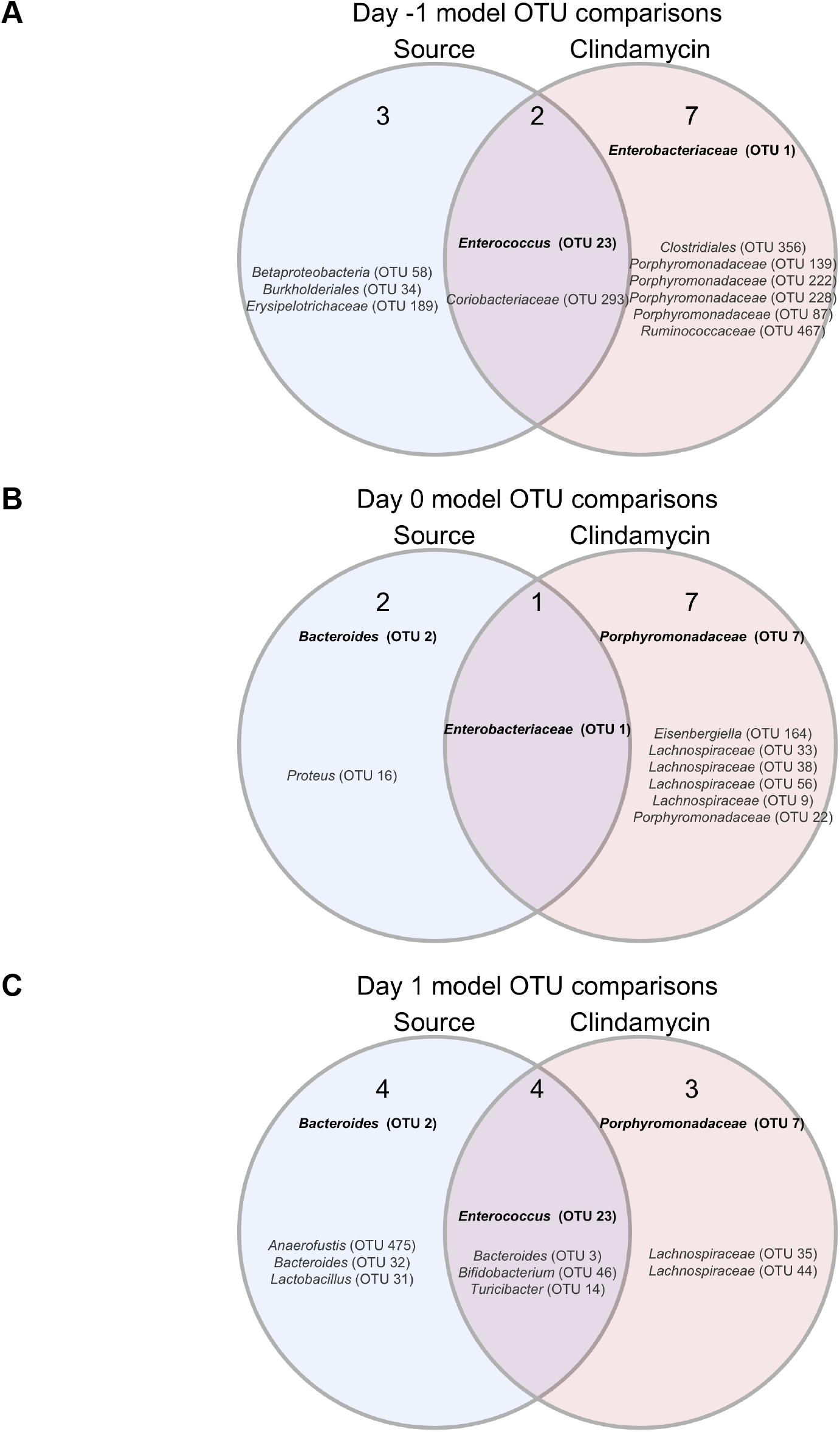
OTUs from classification models based on baseline, post-clindamycin treatment, or 1-day post-infection community data vary by source, clindamycin treatment, or both. A-C. Venn diagrams of OTUs from the top 20 OTUs from the baseline (A), post-clindamycin treatment (B), and 1-day post-infection (C) classification models (Table S14) that overlapped with OTUs that varied across sources at the corresponding timepoint (Tables S5, 8, 10), were impacted by clindamycin treatment (Table S9), or both. Bold OTUs were important to more than 1 classification model.

## Supplementary Tables and Movie

**Movie S1. Large shifts in bacterial community structures occurred after clindamycin and *C. difficile* infection.** PCoA of *θ*_*Y C*_ distances animated from days -1 through 9 of the experiment. Source was the variable that explained the most observed variation across fecal communities (PERMANOVA source R^2^ = 0.35, *P* = 0.0001, Table S11) followed by interactions between cage effects and day of the experiment. Transparency of the symbol corresponds to the day of the experiment, each symbol represents a sample from an individual mouse at a specific timepoint. Circles represent mice from experiment 1 and triangles represent mice from expeirment 2.

**Tables S1-S15. Excel workbook of Tables S1-S15.**

**Table S1. Alpha diversity metrics Kruskal-Wallis statistical results. Table S2. Alpha diversity metrics pairwise Wilcoxon statistical results.**

**Table S3. PERMANOVA results for mice at baseline (day -1), post-clindamycin (day 0), and post-infection (day 1).**

**Table S4. PERMANOVA results for each source of mice at baseline (day -1).**

**Table S5. OTUs with relative abumdances that significantly vary between sources at baseline (day -1).**

**Table S6. *C. difficile* CFU statistical results.**

**Table S7. Mouse weight change statistical results.**

**Table S8. OTUs with relative abundances that significantly vary between sources post-clindamycin (day 0).**

**Table S9. OTUs with relative abundances that significantly changed after clindamycin treatment.**

**Table S10. OTUs with relative abundances that significantly vary between sources 1-day post-infection.**

**Table S11. PERMANOVA results for mice across all timepoints.**

**Table S12. Statistical results of L2-regularized logistic regression model performances compared to random chance.**

**Table S13. Pairwise comparisons of L2-regularized logistic regression model performances.**

**Table S14. Top 20 most important OTUs for each of the 3 L2-regularized logistic regression models based on OTU relative abundance data.**

**Table S15. OTUs with relative abundances that significantly varied between sources of mice on at least 1 day of the experiment by Kruskal-Wallis test.**

